# Spatiotemporal trajectories of formaldehyde fixation effects on quantitative MRI in postmortem human brains

**DOI:** 10.64898/2026.05.05.723107

**Authors:** Yashar Zeighami, Roqaie Moqadam, Liana Sanches, Narges Taghizadehsalehabad, Eve-Marie Frigon, Cecilia Tremblay, Walter Adame-Gonzalez, Dominique Mirault, Zaki Alasmar, Gian Franco Piredda, Gustavo Turecki, Josefina Maranzano, Mallar Chakravarty, Naguib Mechawar, The CIMA-Q group, Mahsa Dadar

## Abstract

**Introduction:** Postmortem human brain magnetic resonance imaging (MRI) offers a unique opportunity to study finer neuroanatomical details and enables direct correlations with gold standard histological and immunohistochemical assessments. However, to prevent tissue decay, postmortem brains are preserved in fixative solutions which can alter tissue properties and exert substantial impacts on the MRI signals. The present study investigates the impact of formalin fixation, the most commonly used solution for postmortem human brain preservation, on different quantitative MRI contrasts.

**Methods:** 142 intact human brain hemispheres immersed in 10% formalin for a range of fixation durations (between 0 days and 20 years) were imaged in a 3T MRI scanner. A subset of 10 brains were further scanned repeatedly at days 0, 3, 10, 20, 30, 60, 90, and 120 to allow for better characterization of the initial transient effects of fixation. Voxel-wise T1 and T2* relaxation, T1/T2 ratio, and myelin water fraction (MWF) maps were generated for each specimen and timepoint, and linear and nonlinear models were used to examine the spatiotemporal changes associated with progressive fixation.

**Results:** All investigated metrics were significantly impacted by formalin fixation, albeit at different rates and with differing regional patterns. T1 and T2* relaxation time decreased as a result of progressive fixation, whereas T1/T2 ratio and MWF measures increased. T1 relaxation and T1/T2 ratio showed nonlinear patterns with initially accelerated changes that decelerate in the first few months, whereas T2* relaxation and MWF changes followed a more linear trend.

**Conclusion:** Formaldehyde fixation exerts systematic changes on quantitative MRI signals that can be modeled and adjusted for to allow for harmonized comparisons of MRI metrics across brains fixed for differing durations. The distinct temporal trajectories observed across metrics highlight the need to account for fixation duration in study design and downstream analyses, particularly when integrating datasets acquired under heterogeneous conditions. Our findings provide a quantitative framework for correcting fixation-induced biases, thereby improving the interpretability and reproducibility of postmortem MRI studies.

## Introduction

Magnetic resonance imaging (MRI) provides a feasible and sensitive approach for studying the human brain. Structural MRI sequences allow for high resolution characterization of brain morphometry and presence of pathology, while quantitative MRI (qMRI) sequences provide robust quantitative estimates of its biophysical tissue properties. In recent years, qMRI studies have shown great promise as biomarkers of different pathologies linked to neurodegeneration, inflammation, and iron accumulation in neurodegenerative disorders (Does, 2018; Parent et al., 2025, 2023; Tuzzi et al., 2020; Weiskopf et al., 2021; Alasmar et al., 2026). However, comprehensive assessments of qMRI metrics as sensitive and specific pathology biomarkers are still lacking. This gap is mainly due to the experimental challenges in performing qMRI sequences and detailed gold standard neuropathological assessments in the same brain tissues.

Postmortem MRI provides a feasible approach to address the translational gap between in vivo imaging and postmortem neuropathology assessments as brain tissues from the same specimens can be imaged and stained consistently to quantify and correlate the presence of specific pathologies (e.g. demyelination, astrogliosis, iron accumulation) across brain disorders (Dadar et al., 2024; Raman et al., 2017). However, major differences between in vivo and ex vivo settings impact the MRI signal in postmortem settings (Dadar et al., 2024; Pfefferbaum et al., 2004). Instead of in situ body temperature (∼37 ℃), ex vivo brain specimens are generally scanned at room temperature (∼21 ℃), and these temperature differences lead to systematic differences in certain MRI metrics (Birkl et al., 2016). Furthermore, to avoid tissue degradation, specimens are generally immersed in a fixative solution, most commonly 10% Neutral buffered formalin (NBF), i.e. 4% formaldehyde, which also significantly impacts the MRI signal (Dadar et al., 2024; Frigon et al., 2025, 2026a; Pfefferbaum et al., 2004).

Longitudinal (T1) and transverse (T2) relaxation times of water protons rely on water content within the tissue and its mobility (Pfefferbaum et al., 2004). Formaldehyde fixation impacts tissue microstructure, reducing water mobility (Shepherd et al., 2009) and leading to reduction in T1 and T2 values (Shatil et al., 2018) which consequently impacts diffusion weighted and myelin water imaging signals (Miller et al., 2011; Roebroeck et al., 2019; Xu et al., 2025). These changes also impact gray-to-white matter (GM-WM) tissue contrast as T1 values in gray matter (GM) and white matter (WM) converge (Pfefferbaum et al., 2004; Tovi and Ericsson, 1992). Furthermore, since tissues at the outer surface of the brain are exposed to the fixative first compared to deeper tissues while brains are immersed in NBF, these alterations are not consistent across all brain regions (Raman et al., 2017), potentially impacting downstream MRI based quantifications (Neuhaus et al., 2025). Based on prior literature, these effects are mainly observed in the first weeks of fixation, and T1 and T2 values reportedly stabilize afterwards (Pfefferbaum et al., 2004), though rates of stabilization can differ based on multiple factors including tissue type and its proximity to surface (Raman et al., 2017). However, the exact patterns of fixation and stabilization and whether these patterns differ across regions and tissue types have not been established.

While scanning all postmortem specimens at the same temperature (i.e. ∼21 ℃) effectively minimizes the impact of temperature variability on the ex vivo MRI signals, to reach adequate sample sizes that represent the characteristics of the groups or pathologies of interest, most postmortem MRI studies inevitably integrate data from samples fixed with a wide range of fixation durations (Dawe et al., 2011, 2009; Duijn, 2018; Frigon et al., 2025; Khandelwal et al., 2024; Laule et al., 2006; McAleese et al., 2021, 2017; Neuhaus et al., 2025; Roseborough et al., 2020). Given that formaldehyde fixation has nonnegligible impacts on certain qMRI metrics, it is necessary to model the effect of fixation duration on qMRI signals to enable systematic adjustments that account for these effects in downstream analyses.

The present study addresses this question by quantifying the impact of prolonged formaldehyde fixation on qMRI metrics. Using an unprecedented sample of 140 postmortem brains fixed between 0 days to 20 years with a diverse range of neurodegenerative pathologies, we examined the impact of formaldehyde fixation on T1 and T2* relaxation times, T1/T2 ratio, and myelin water fraction (MWF) across different tissue types, regions, and distances from the tissue surface.

## Methods

Ethics approval for the study was obtained from the Ethics Review Board of the Douglas Research Center (2023-622). Postmortem human brain specimens were acquired from the Douglas Brain Bank (DBB), the largest brain bank in Canada housing over 3600 brain specimens with different neurodegenerative disorders including Alzheimer’s disease (AD), Parkinson’s disease (PD), frontotemporal dementia (FTD), vascular dementia (VD), Lewy body dementia (LBD), Amyotrophic lateral sclerosis (ALS), as well as diverse psychiatric conditions including schizophrenia, major depression, bipolar disorder, and substance use disorders (https://douglasbrainbank.ca/). Brain specimens are collected in accordance with informed consent of the donors or their next of kin, according to tissue banking practices regulated by the Quebec Health Research Fund, and the Guidelines on Human Biobanks and Genetic Research Databases overseen by the Douglas Research Ethics Board (Dadar et al., 2024).

A subset of the specimens (N = 10) were from participants in the Consortium for the Early Identification of Alzheimer’s Disease-Quebec (CIMA-Q) (Belleville et al., 2019). The CIMA-Q is a multicenter observational study that includes a wide clinical spectrum of aging and AD across multiple sites in Quebec. Participants with normal cognition, subjective cognitive decline, mild cognitive impairment (MCI), and mild AD dementia were recruited through memory clinics, community ads, and pre-existing research cohorts. Eligibility for the CIMA-Q study required participants to be aged 65 years or older, able to complete clinical and cognitive assessments, proficient in French or English, and have no contraindications to MRI.

Following reception at the DBB, hemispheres are separated by a midline sagittal cut through the brain, brainstem, and cerebellum. One intact hemisphere (right or left, in alternation) is fixed by immersion in 6 liters of 10% formalin. After 3 weeks of fixation, specimens are transferred to and remain in air-tight MRI-compatible plastic containers fully immersed in the same 10% NBF. MRI acquisitions were performed in the same containers, to minimize tissue manipulation and prevent exposure of tissue to air, leading to inevitable MRI artifacts. To study the effect of formaldehyde fixation on MRI signals during the first 3-week period, specimens were transferred from the 7-liter containers to the same containers that are used for long term storage, and were consistently scanned in the same containers (Dadar et al., 2024).

A total of 142 aging specimens including cases without neurological disorders as well as specimens with clinical and neuropathological diagnoses of neurodegenerative disorders (AD, VD, PD, LBD, FTD, and ALS) spanning a wide range on variables of interest (i.e., formaldehyde fixation time, PMI, and pH) were included. To enable assessment of intra specimen changes and variabilities while maintaining feasibility, a subset of 10 specimens spanning all diagnostic groups were scanned consistently at 8 fixation timepoints including 0, 3, 10, 20, 30, 60, 90, and 120 days following death. Furthermore, a second subset of 10 specimens covering all diagnostic groups were scanned twice with an approximate one year fixation interval. The rest of the specimens were scanned once, with fixation times ranging between 0 days and 20 years. Table 1 summarizes the characteristics of the postmortem specimens included in this study.

**Table 1.** Characteristics of the specimens used in this study.

|  | **Aging** | **AD/VD** | **ALS/FTD** | **PD/LBD** | **Other** | **All** |
| --- | --- | --- | --- | --- | --- | --- |
| **N** | 13 | 56 | 17 | 24 | 32 | 142 |
| **N<sub>Female</sub> (%)** | 4 (30%) | 25 (45%) | 6 (35%) | 9 (37%) | 17 (53%) | 61 (42%) |
| **N<sub>Timepoint</sub><sup>1</sup>** | 32 | 89 | 39 | 35 | 47 | 242 |
| **Age at death (years)** | 79.4 ± 14.0 | 83.5 ± 8.9 | 68.1 ± 12.3 | 76.7 ± 8.2 | 73.6 ± 10.9 | 79.9 ± 11.1 |
| **Brain Weight (kg)** | 1.2 ± 0.22 | 1.09 ± 0.13 | 1.08 ± 0.23 | 1.16 ± 0.14 | 1.17 ± 0.17 | 1.11 ± 0.15 |
| **PMI (hours)** | 31.3 ± 24.1 | 28.5 ± 14.9 | 26.9 ± 15.6 | 21.7 ± 14.9 | 29.7 ± 19.1 | 26.7 ± 15.2 |
| **Fixation Time (years)** | 4.1 ± 4.7 | 12.8 ± 6.3 | 5.8 ± 6.7 | 9.4 ± 6.4 | 7.3 ± 6.2 | 10.64 ± 6.8 |
| **pH** | 6.07 ± 0.39 | 5.87 ± 0.20 | 6.05 ± 0.20 | 6.1 ± 0.26 | 6.1 ± 0.29 | 6.00 ± 0.29 |

AD: Alzheimer's dementia. VD: Vascular Dementia. PD: Parkinson's Disease. LBD: Lewy Body Dementia. ALS: Amyotrophic Lateral Sclerosis. FTD: Frontotemporal Dementia. PMI: Postmortem interval.
1. Timepoint reflects the number of times the specimens have been scanned to examine the effect of fixation over time. 2. Other diagnoses include multiple systems atrophy (MSA), progressive supranuclear palsy (PSP), corticobasal degeneration (CBD), and hippocampal sclerosis

MRI acquisitions were performed using a 3T Siemens Prisma_Fit human MRI scanner at the Douglas Cerebral Imaging Centre (CIC). The containers were stabilized in a 64-channel head/neck coil, with the hemispheres lying flat on the sagittal cut in the center of the container (i.e. not touching the sides) and the cerebellum towards in-bore to standardize positioning and minimize artifacts. Structural T1-weighted (T1w) and T2-space MRIs were acquired for tissue segmentation and deriving T1/T2 ratio maps. Three 3D multi-echo gradient echo (meGRE) sequences were acquired with different flip angles to calculate quantitative T1 and T2* maps. A 3D multi-echo gradient and spin echo (GRASE) research application sequence was also acquired to derive T2 and myelin water fraction (MWF) maps. Table 2 summarizes the acquisition parameters for these sequences.

**Table 2.**
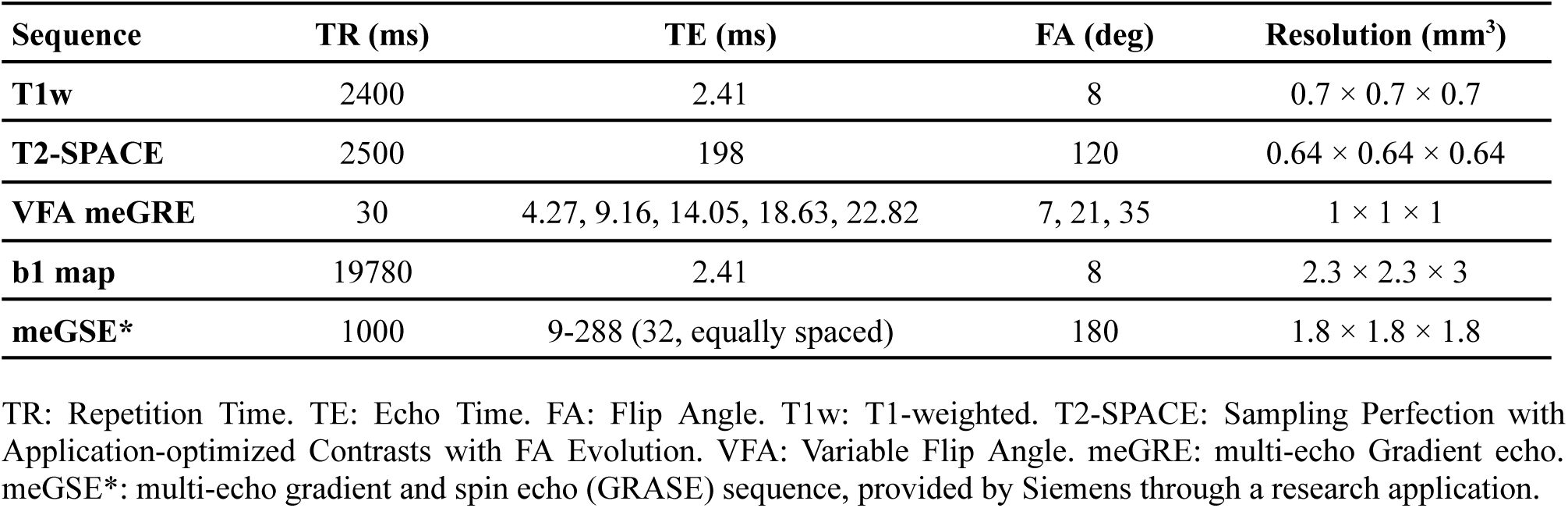
Acquisition protocol for the sequences used in this study.

Figure 1 summarizes the image processing framework. Structural T1w and T2w images were processed using an in-house pipeline (https://github.com/VANDAlab/Ex_vivo_Pipeline) built on open source and open access tools. Preprocessing steps included denoising (Coupé et al., 2008), intensity inhomogeneity correction (Sled et al., 1998), and intensity normalization into range 0-100.

**Figure 1.**
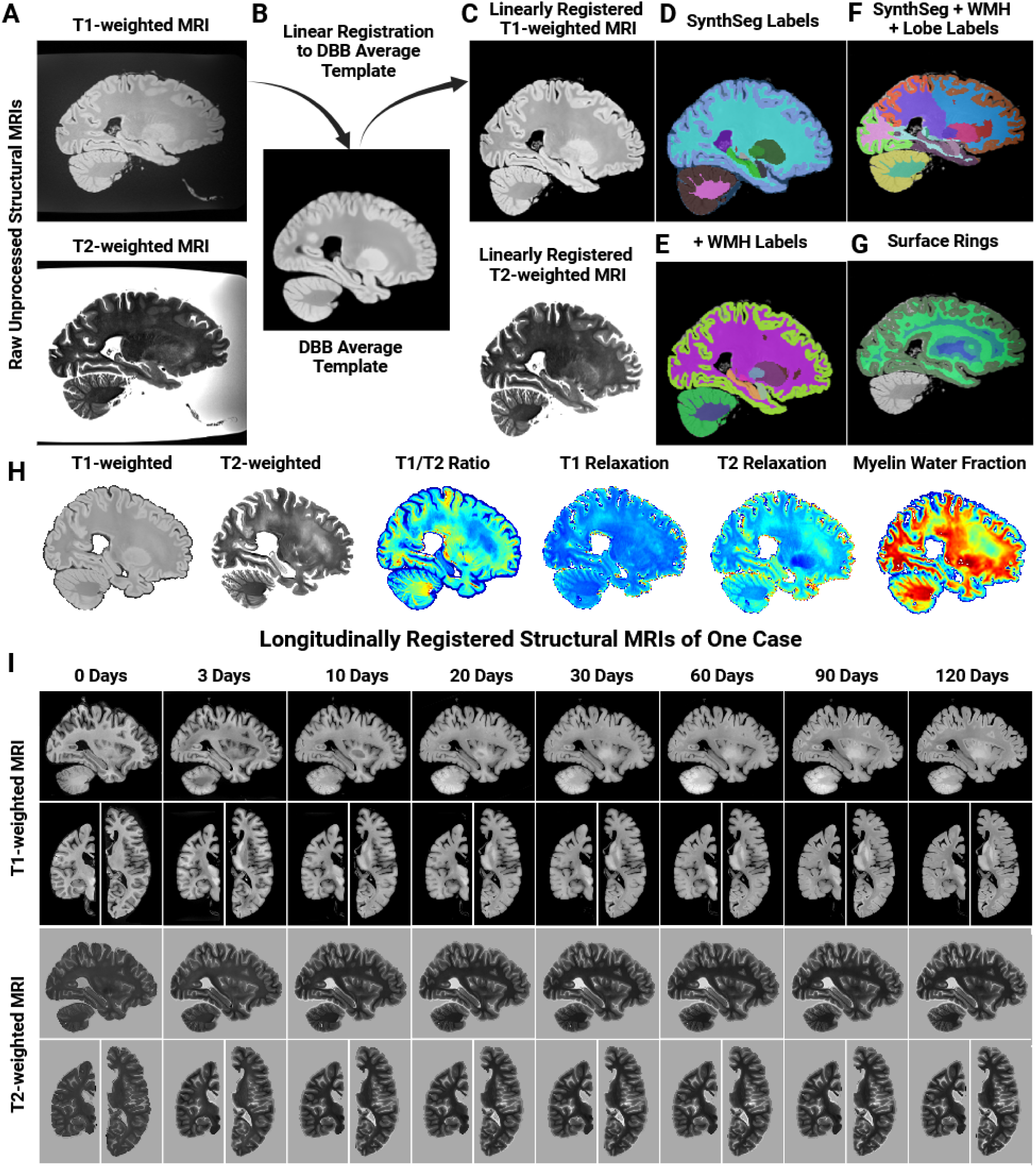
Structural MRI processing framework diagram. **A**. Raw T1-weighted and T2-weighted MRIs. **B**. Linear registration to DBB average template. **C**. Preprocessed T1-weighted and T2-weighted images are linearly registered to the DBB average template in the stereotaxic space. **D**. SynthSeg is applied to the linearly registered and preprocessed T1-weighted images to derive an initial mask. **E-F**. WMH and lobar labels are added to the SynthSeg mask. **G**. Surface rings are derived based on the brain mask. **H**. Examples of derived structural and quantitative MRI maps for one DBB specimen. **I**. Longitudinally registered structural T1-weighted and T2-weighted MRIs of one DBB specimen scanned at 0, 3, 10, 20, 30, 60, 90, and 120 days, showing the impact of formaldehyde fixation on signal and contrast changes in structural sequences. DBB: Douglas Brain Bank.

The preprocessed images were then linearly registered to the stereotaxic space using an in-house ex vivo template derived based on our ex vivo T1w images (Dadar et al., 2018a, 2024). SynthSeg was applied to the preprocessed T1w images in the stereotaxic space for tissue segmentation (Figure 1, Panel D) (Iglesias et al., 2023). Left and right hemisphere labels were merged and white matter hyperintensities (WMH) were manually segmented and added. Hammers lobar atlas was used to derive lobar cortical gray matter (cGM) and WM segmentations (Dadar et al., 2018b).

Tissue masks were transformed back to the native space to extract qMRI signals in the native resolution without resampling. These masks were also used to generate depth ring masks distanced at 2mm intervals from the surface by using a morphological erosion operation to examine the rate of fixation in superficial versus deep brain tissues. For the longitudinally scanned specimens, the last timepoint (with the highest GM-WM tissue contrast) was used for tissue segmentation, and the resulting labels were registered to all other timepoints based on rigid 6-parameter registrations to minimize the impact of segmentation related variability across timepoints (Figure 1, Panel I). To examine the impact of formaldehyde fixation duration on signal to noise ratio (SNR), mean SNR values were calculated both inside each brain region of interest (ROI) as well as inside 5 manually segmented cylindrical masks with a radius of 12 voxels and height of 10 voxels (i.e. a volume of ∼4500 voxels) in the NBF solution in each corner as well as the center of the container.

qMRLab was used to generate quantitative T1 and T2* maps based on the VFA meGRE acquisitions as well as MWF maps based on meGSE acquisitions (Duval et al., 2018; Karakuzu et al., 2019). T1/T2 ratio maps were derived following linear co-registration of the T1w and T2w images in the stereotaxic space. Median T1 and T2* relaxation, T1/T2 ratio, and MWF values were extracted for each specimen and label. All image processing steps were visually assessed to ensure the quality of the derived measures, and cases that did not pass this visual quality control step were excluded from subsequent analyses.

### Statistical Analyses

Linear and nonlinear models were used to characterize the relationship between post-mortem fixation duration and qMRI measures (T1 relaxation, T2* relaxation, MWF, and T1/T2 ratio). In each case, voxel-wise qMRI maps were summarized at regional level using median values, with zero-valued voxels removed. Analyses were conducted across cGM, WM, WMH, and dGM regions, as well as across concentric brain depth “rings” for WM and GM compartments within each cortical division.

In all cases, linear models were used as the base model, and other candidate models were compared to the base linear model. Model selection was performed using the Bayesian Information Criterion (BIC), allowing comparison across models with differing complexity.

#### T1 relaxation

Alternative nonlinear regression models, including exponential and power-law formulations, were fitted per region to describe the dependence of T1 relaxation on fixation time. Model parameters were estimated using nonlinear least squares (Levenberg–Marquardt algorithm), and standard errors, Wald confidence intervals, and p-values were derived from the fitted models. Based on these comparisons, a power-law decay model (formula I) was selected as the primary model, as it provided the best fit across regions. Residuals from the fitted models were extracted to obtain fixation-adjusted T1 measures for downstream analyses.

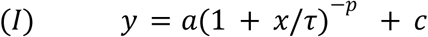

#### T1/T2 Ratio

A nonlinear shifted-logarithmic model was assessed as an alternative to the base linear model to better capture the observed temporal dynamics. Shifted-log models allow for rapid changes at short fixation times while remaining well-defined at *x* = 0 through the inclusion of a positive shift parameter (*x*_0_).

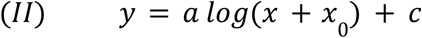

Model parameters were estimated using nonlinear least squares, and inference was performed as described in the previous section. Residuals from the selected models were computed to obtain fixation-adjusted T1/T2 ratio measures. *a* is a scaling parameter governing the long-term rate of change, *x*_0_ is a positive shift parameter controlling early-time curvature, and *c* is a baseline offset.

To ensure numerical stability and enforce the constraint *x*_0_ > 0, the shift parameter was reparameterized as *x*_0_ = *exp(lx*_0_), and the model was fit in terms of *lx*_0_ The long-term rate of change is governed by *a* and the early-time value at *x* = 0 initial slope at fixation onset is given by *a*/*x*_0_.

#### T2* and MWF

A linear modeling framework was adopted for T2* and MWF measures, consistent with the approaches used in the previous analyses. Specifically, regional values were modeled as a function of fixation time, and residuals were extracted to obtain fixation-adjusted measures.

All model fits were performed independently for each brain region, and fixation-adjusted residuals were used to normalize WMH measures relative to corresponding WM regions, thereby accounting for shared fixation-related variance. False discovery rate (FDR) correction was used to account for multiple comparisons across regions and parameters (Benjamini and Yekutieli, 2005). For residual analyses, Holm correction was used to account for multiple comparisons.

In order to directly compare trajectories between regions, a set of nested nonlinear (and linear) models with increasing levels of regional specificity were fitted. In each case, a shared model (common parameters across regions) was compared to a “full region-specific model” (in which all parameters were allowed to vary by region). In addition, a “shape model” was defined in which only parameters governing the trajectory shape were allowed to vary across regions, while baseline terms were held constant. For T2* and MWF, a linear model was used, and the shape model allowed regional variation in the slope parameter while the intercept was shared. For T1 relaxation, a power-law model was used, and the shape model allowed regional variation in the timescale (*τ*) and exponent (*p*) parameters, while (*a*) and (*c*) were shared. For T1/T2 ratio, a shifted-log model was used, and the shape model allowed regional variation in (*a*) and (*x*_0_), while (*c*) was shared.

Model fit was evaluated using the BIC. For all models including regional effects, the occipital cortex was used as the reference region for cGM, WM, and WMHs, while pallidum was used as the reference region for dGM structures.

### Data and Code Availability Statement

The ex vivo image processing pipeline and qMRI extraction scripts are available at https://github.com/VANDAlab/Ex_vivo_Pipeline. Data from the CIMA-Q participants is open access and can be requested via the CIMA-Q platform (https://www.cima-q.ca).

## Results

### T1 relaxation changes associated with formaldehyde fixation

Figure 2 shows the changes in T1 relaxation times associated with formaldehyde fixation for cGM and dGM (Panels A and B respectively), as well as WM, and WMHs (Panels C and D). Panels E-G show the same trajectories for a longitudinally scanned specimen during its first four months of fixation. Panel H shows longitudinally registered axial, coronal, and sagittal views of T1 relaxation maps of the same specimen across its longitudinal timepoints. Across all regions, T1 relaxation times exhibited a rapid decline during the first weeks of fixation followed by gradual stabilization over longer time periods.

**Figure 2.**
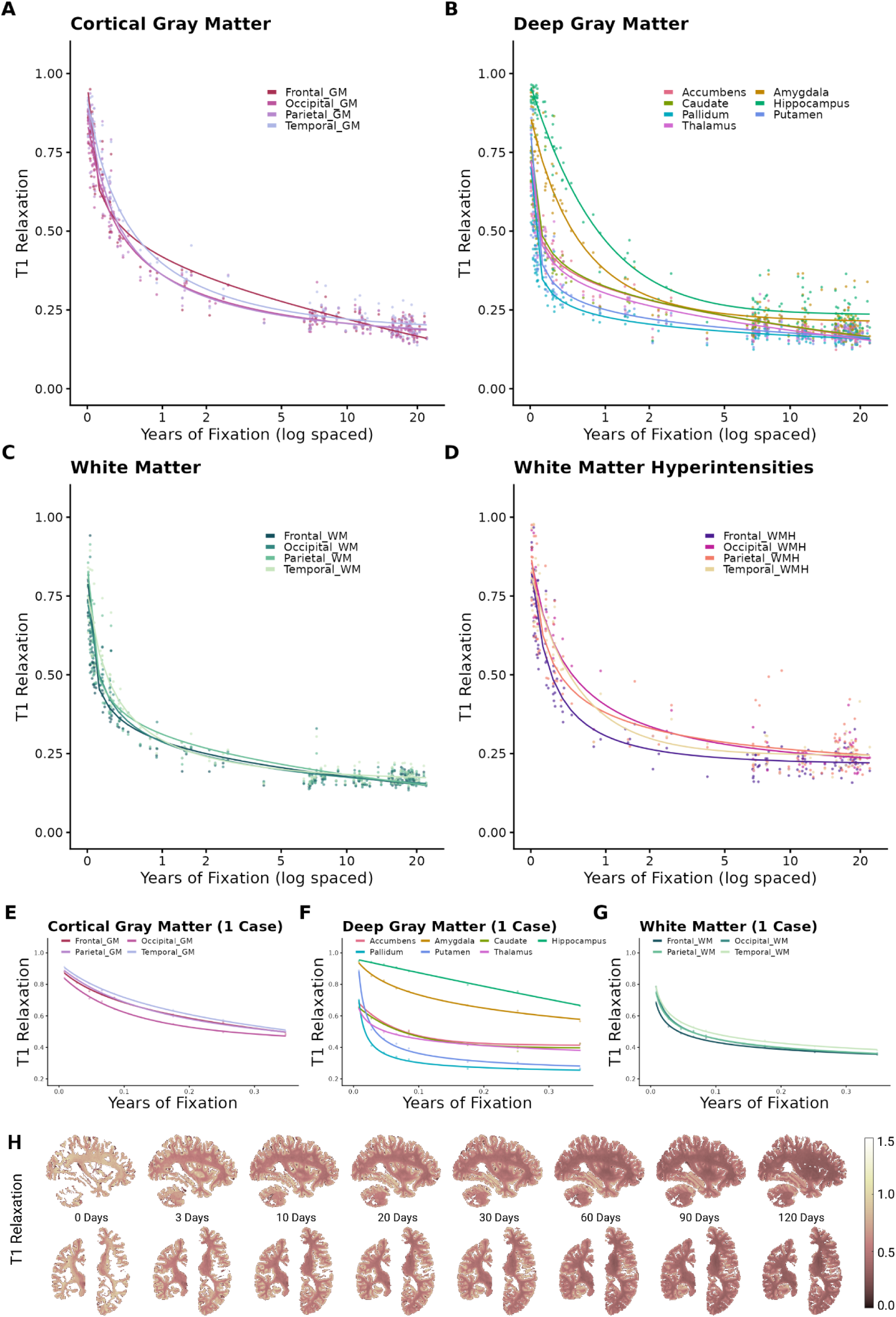
T1 relaxation changes associated with formaldehyde fixation. **A.** Lobar cortical GM regions. **B.** Deep GM regions. **C.** Lobar WM regions. **D.** Lobar WMH regions. **E-G.** Changes in lobar cortical GM, deep GM, and WM regions in one hemisphere scanned longitudinally from 0 to 120 days of fixation. **H.** Sagittal, coronal, and axial slices showing voxel-wise T1 relaxation maps of the same longitudinally scanned specimen from 0 to 120 days of fixation. GM: Gray Matter. WM: White Matter. WMH: White Matter Hyperintensity.

cGM trajectories showed clear regional differences, with faster decline and earlier stabilization in the occipital cortex, followed by frontal and parietal regions, and slower trajectories in the temporal cortex. These qualitative patterns were confirmed by model-based analyses, where the power-law decay model outperformed linear and exponential alternatives. Model comparison using BIC indicated the best fit by the shape model compared to both shared and full region-specific models (BIC: −2090, −2041, and −2066, respectively), suggesting regional variability in trajectory shape rather than baseline shifts. Relative to the occipital cortex (τ=0.10), τ was significantly increased in the parietal (β = 0.055, p = 0.009) and temporal cortex (β = 0.126, p < 0.001), indicating slower decay in these regions, while no significant difference was observed in the frontal cortex (p = 0.25).

Similarly, in WM, trajectories exhibited rapid early declines followed by stabilization, but with less pronounced regional differentiation. The shape model outperformed both shared and full region-specific models (BIC: −1990, −1960, and −1952, respectively). Relative to the occipital cortex (τ=0.01), τ was significantly increased in the temporal lobe (β = 0.009, p < 0.001), indicating slower decay. For WMH, while the shape model outperformed the other alternatives, there were no statistically significant regional differences in either τ or p parameters.

In contrast, for dGM regions, the full region-specific model provided the best fit (BIC = −3139), outperforming both the shared and shape models (BIC = −1735 and −3129, respectively), indicating that regional variability extends beyond differences in trajectory shape alone. Parameter estimates demonstrated widespread regional heterogeneity across all model components. Significant differences were observed in the timescale parameter (τ), with increases in the hippocampus (β = 1.49, p = 0.037) and amygdala (β = 0.48, p = 0.019), and a decrease in the accumbens (β = −0.005, p = 0.044), relative to the pallidum (τ=0.008). In addition, robust regional differences were observed in the exponent parameter (p), with reductions in the thalamus (β = −0.29, p < 0.001), accumbens (β = −0.32, p < 0.001), and caudate (β = −0.26, p < 0.001), indicating substantial variation in trajectory curvature across regions. Regional differences were also observed in the baseline levels (c), with increases in the hippocampus (β = 0.090, p < 0.001) and amygdala (β = 0.062, p < 0.001), and decreases in the thalamus (β = −0.152, p = 0.008), accumbens (β = −0.210, p = 0.031), and caudate (β = −0.125, p = 0.022). Similarly, scaling parameter (a) varied significantly across regions, with increases in the thalamus (β = 0.19, p = 0.005), accumbens (β = 0.22, p = 0.039), and caudate (β = 0.16, p = 0.021), and a decrease in the amygdala (β = −0.087, p = 0.013).

Together, these findings indicate a differential pattern of regional variability in fixation-related dynamics, with strong and shape-driven differences in cGM, reduced and more localized effects in WM, and highly heterogeneous, multi-parameter differences in dGM structures.

### T2* relaxation changes associated with formaldehyde fixation

Figure 3 illustrates the changes in T2* relaxation times for cGM and dGM (Panels A and B respectively), WM, and WMHs (Panels C and D). Similarly, Panels E-G and H show longitudinal trajectories and voxel-wise T2* relaxation maps of the same specimen across its longitudinal timepoints. Overall, T2* relaxation times exhibited more subtle and approximately linear changes over fixation time compared to T1, with region-dependent differences in both baseline levels and rates of change.

**Figure 3.**
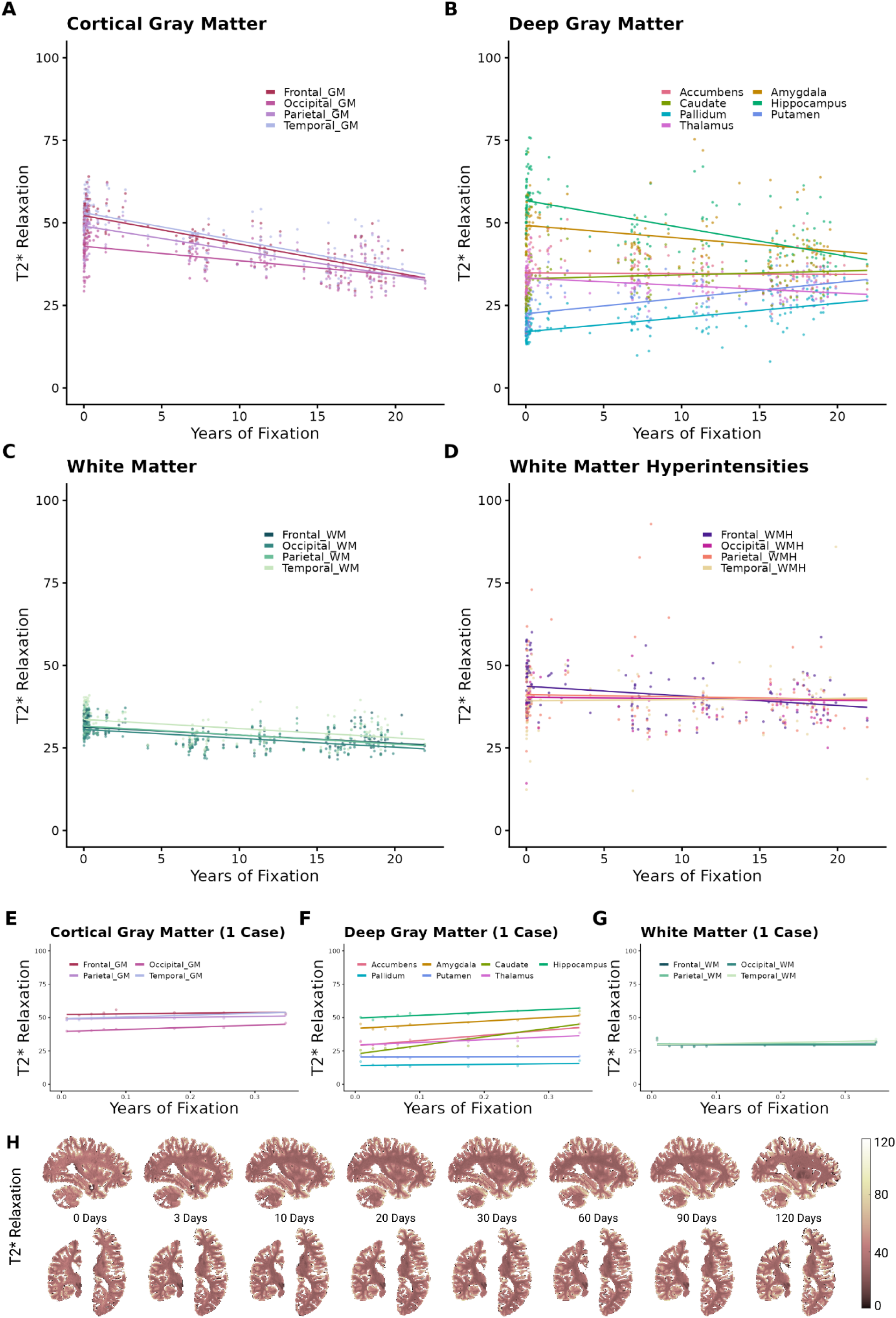
T2* relaxation changes associated with formaldehyde fixation. **A.** Lobar cortical GM regions. **B.** Deep GM regions. **C.** Lobar WM regions. **D.** Lobar WMH regions. **E-G.** Changes in lobar cortical GM, deep GM, and WM regions in one hemisphere scanned longitudinally from 0 to 120 days of fixation. **H.** Sagittal, coronal, and axial slices showing voxel-wise T2* relaxation maps of the same longitudinally scanned specimen from 0 to 120 days of fixation. GM: Gray Matter. WM: White Matter. WMH: White Matter Hyperintensity.

Within cGM, model comparison using BIC indicated that the full region-specific model provided the best fit compared to both shared and shape models (BIC = 4126, 4281, 4280, respectively). The occipital cortex showed a significant negative slope (β = −0.42, p < 0.001) indicating a decline in T2* values over time. The frontal, parietal, and temporal cortices exhibited significantly higher baseline values relative to occipital cortex (β = 9.06, 6.00, 9.89, respectively, all p-values < 0.001), along with significantly steeper negative slopes (β = −0.41, −0.30, −0.40, respectively, all p-values < 0.001), indicating faster decline over time compared to occipital lobe (Figure 3-A). Similarly, in the WM, model comparison using BIC indicated that the full region-specific model provided the best fit compared to both shared and shape models (BIC = 3868, 3873, 3874, respectively). The occipital WM showed a significant negative slope (β = −0.34, p < 0.001) indicating a decline in T2* values over time, while no significant regional differences in slopes were observed, indicating largely homogeneous rates of change across WM regions. The temporal WM exhibited significantly higher baseline values relative to occipital cortex (β = 3.17, p < 0.001) (Figure 3-C).

WMH trajectories demonstrated minimal changes and no meaningful regional variation. Model comparison favored the shared model (BIC = 3627) over both the shape and full region-specific models (BIC = 3645, 3658, respectively), indicating that a single common trajectory adequately describes T2* changes across WMH regions. This is consistent with the small but significant global decline observed over time (β = −0.283, p < 0.001), without evidence for regional effects (Figure 3-D).

In contrast, dGM exhibited marked heterogeneity in fixation-related T2* changes. Model comparison strongly favored the full region-specific model (BIC = 7542) over both shared and shape models (BIC = 8848 and 8703), indicating substantial variation in both baseline levels and rates of change. Baseline T2* values differed markedly across structures, with particularly higher values in hippocampus (β = 39.25, p < 0.001) and amygdala (β = 32.24, p < 0.001), as well as in thalamus, accumbens, and caudate (all p < 0.001). In addition, slopes varied significantly across regions, with hippocampus (β = −1.22, p < 0.001), amygdala (β = −0.82, p < 0.001), thalamus (β = −0.69, p < 0.001), accumbens (β = −0.43, p < 0.001), and caudate (β = −0.25, p = 0.012) showing steeper declines relative to the pallidum (reference region), while putamen showed no significant difference.

### T1/T2 ratio changes associated with formaldehyde fixation

Figure 4 shows the changes in T1/T2 ratios for cGM, dGM (Panels A and B respectively), WM, and WMH (Panels C and D) regions and longitudinal trajectories (Panels E-G) and voxel-wise maps (Panel H). While shifted-logarithmic model still showed a better fit due to the initial increase in the first days, T1/T2 ratio changes in the cGM generally followed a linear trajectory, with the occipital lobe showing the steepest slope, followed by parietal, temporal, and frontal cGM. dGM, WM, and WMH trajectories showed a more pronounced logarithmic trend, with an initial increase period in the first few weeks, followed by a linear increase after the first year of fixation.

**Figure 4.**
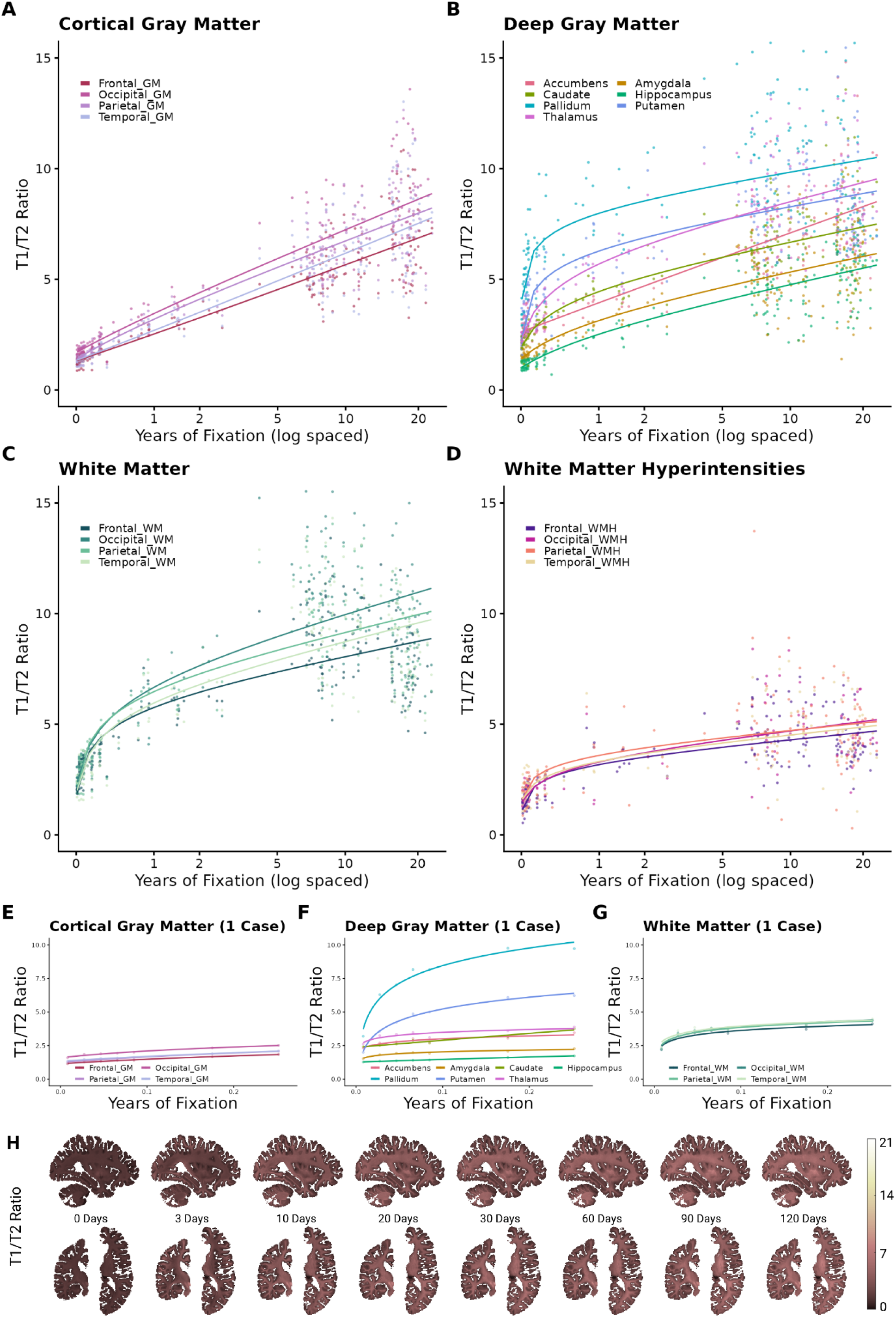
T1/T2 ratio changes associated with formaldehyde fixation. **A.** Lobar cortical GM regions. **B.** Deep GM regions. **C.** Lobar WM regions. **D.** Lobar WMH regions. **E-G.** Changes in lobar cortical GM, deep GM, and WM regions in one hemisphere scanned longitudinally from 0 to 120 days of fixation. **H.** Sagittal, coronal, and axial slices showing voxel-wise T1/T2 ratio maps of the same longitudinally scanned specimen from 0 to 120 days of fixation.

Within cGM, model comparison using BIC indicated that the shape model provided the best fit (BIC = 2572), outperforming both the shared (BIC = 2641) and full region-specific models (BIC = 2590). This indicates that regional variability was primarily driven by differences in trajectory shape rather than baseline shifts. Relative to occipital cortex (*a*=2.22), all regions showed significantly lower scaling parameters (all p < 0.01), with additional significant differences in early-trajectory curvature in the frontal and temporal cortices, while initial slopes were highly similar across cGM regions. In WM, the shape model also provided the best fit (BIC = 3143), outperforming both the shared (BIC = 3165) and full models (BIC = 3149). Regional differences were evident in both scaling and early-trajectory curvature, with frontal, parietal, and temporal WM showing significantly lower values relative to occipital WM (all p < 0.001).

For WMH, the shared model provided the best fit (BIC = 1798), outperforming both the shape (BIC = 1831) and full models (BIC = 1843). This indicates that a common shifted-log trajectory adequately described T1/T2 ratio changes across WMH regions, with no evidence that regional parameter differences improved model fit. The shared model showed a significant positive scaling parameter (*a* = 0.50, p < 0.001), consistent with increasing T1/T2 ratio over fixation time followed by stabilization.

In contrast, dGM showed substantial regional heterogeneity. Model comparison strongly favored the fully region-specific model (BIC = 5050), outperforming both the shared (BIC = 5848) and shape models (BIC = 5268), indicating that regional variability extended beyond shape differences alone. Relative to pallidum, thalamus and accumbens showed significantly increased long-term scaling, with *a* increasing from 0.82 in pallidum to 1.19 in thalamus and 1.64 in accumbens. However, the derived initial slopes revealed a different pattern: pallidum showed a steep initial slope, followed by putamen, thalamus, and caudate (21.38), while amygdala, hippocampus, and accumbens showed much slower initial rates. Early values were also heterogeneous, ranging from 0.97 in hippocampus to 2.25 in accumbens. These findings indicate that dGM regions differ not only in baseline levels and long-term scaling, but also in the timing and sharpness of early fixation-related changes.

Together, these findings indicate that T1/T2 ratio trajectories follow a shifted-log pattern across regions, with shape-driven differences in cGM and WM, minimal regional differentiation in WMH, and pronounced multi-parameter heterogeneity in dGM. More specifically, cGM regional differences were dominated by long-term scaling and early offset and dGM differences reflected combined changes in early slope, early offset, long-term scaling, and baseline level.

### MWF changes associated with formaldehyde fixation

Figure 5 shows the changes in MWF for cGM, dGM, WM, and WMH regions (Panels A-D, respectively), longitudinal trajectories (Panels E-G), and voxel-wise MWF maps (Panel H). Across tissue classes, MWF values showed an approximately linear increase with fixation time.

**Figure 5.**
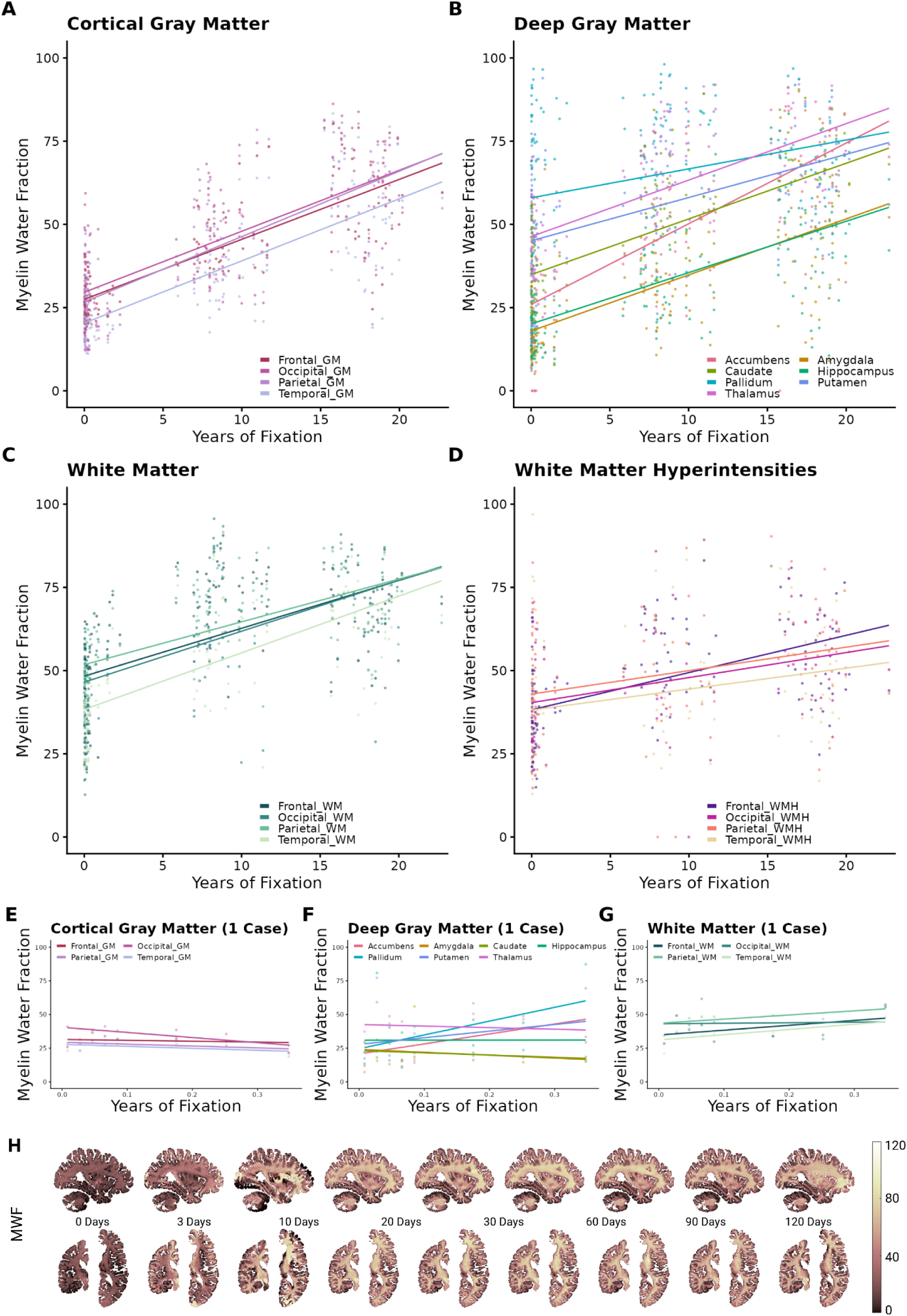
MWF changes associated with formaldehyde fixation. **A.** Lobar cortical GM regions. **B.** Deep GM regions. **C.** Lobar WM regions. **D.** Lobar WMH regions. **E-G.** Changes in lobar cortical GM, deep GM, and WM regions in one hemisphere scanned longitudinally from 0 to 120 days of fixation. **H.** Sagittal, coronal, and axial slices showing voxel-wise MWF maps of the same longitudinally scanned specimen from 0 to 120 days of fixation. MWF: Myelin Water Fraction. GM: Gray Matter. WM: White Matter. WMH: White Matter Hyperintensity.

Within cGM, model comparison using BIC indicated that the full region-specific model provided the best fit (BIC = 5052), outperforming both the shared (BIC = 5064) and shape models (BIC = 5057). However, this improvement was primarily driven by baseline differences rather than slope differences, as regional slope terms were not significant. Relative to occipital cortex, temporal cortex showed a significantly lower baseline MWF (β = −8.83, p < 0.001), while frontal and parietal cortices did not differ significantly. The occipital lobe slope was positive (β = 1.92, p < 0.001), indicating increasing MWF with fixation time across cGM regions, with other regions not showing any statistically significant slope differences.

Similarly, in WM, the full region-specific model provided the best fit (BIC = 5319), although the difference from the shared model was modest (BIC = 5324), with no statistically significant regional slope difference compared to that of occipital lobe, which significantly increased (β = 1.72, p < 0.001). Baseline MWF differed across WM regions, with higher values in parietal WM (β = 4.57, p = 0.045) and lower values in temporal WM (β = −7.86, p < 0.001) relative to occipital WM. For WMH, the shared model provided the best fit (BIC = 3776), outperforming both the shape (BIC = 3786) and full region-specific models (BIC = 3800). This indicates that a common trajectory adequately describes MWF changes across WMH regions. The shared model showed an intercept of β = 37 with a significant positive slope (β = 1.04, p < 0.001).

In contrast, dGM showed substantial regional heterogeneity. Model comparison strongly favored the full region-specific model (BIC = 9152), outperforming both the shared (BIC = 9524) and shape models (BIC = 9387). Relative to pallidum (β = 54.5), all dGM regions showed significantly lower baseline MWF values, with the largest differences observed in amygdala (β = −37.27, p < 0.001) and hippocampus (β = −35.49, p < 0.001). Slopes also differed significantly across structures, with faster increases in thalamus (β = 0.80, p = 0.002), hippocampus (β = 0.51, p = 0.044), amygdala (β = 0.63, p = 0.016), accumbens (β = 1.33, p < 0.001), and caudate (β = 0.69, p = 0.007) relative to that of pallidum (β = 1.1, p < 0.001), while putamen did not differ significantly.

Together, these findings indicate that fixation-related MWF changes are characterized by a global linear increase over time, with little evidence for regional slope differences in cGM, WM, or WMH, but pronounced regional heterogeneity in dGM, where both baseline levels and rates of increase differed across structures.

### Tissue and depth related differences in fixation patterns

To examine whether potential differences in penetration of formaldehyde impact the rates of change in T1 and T2* relaxation, T1/T2 ratio, and MWF, these measures were compared across different depths (see Figure 1. Panel G. Surface Rings) of cortical and WM regions (Figure 6). In the first year following fixation, T1 relaxation values declined more slowly in WM and GM regions closer to the surface of the tissue (Figure 6-A, Ring 1) compared to deeper rings, and WM signal declined faster compared to GM (Figure 6-E). This pattern was relatively consistent across lobes (Supplementary Figures 1-3), with the exception of the occipital WM where Ring 3 had a slower (i.e. deep WM) decline compared to Rings 1 and 2. A different pattern was observed for T2* relaxation in the GM, where tissues closer to the surface (Figure 6-B. Ring 1) declined relatively faster. Compared to cortical GM, WM T2* values declined more slowly (Figure 6-F), and did not differ across depths. In the first year of fixation, T1/T2 ratio values increased at slower rates in the rings closer to the surface (Figure 6-D), and WM T1/T2 ratios declined at substantially faster rates during the first year compared to cortical GM, which declined almost linearly (Figure 6-H). Finally, GM MWF values had overall higher slopes closer to the surface (Figure 6-C). WM MWF values increased faster in the WM ring closer to the surface, and did not differ in their rates of increase compared to cortical GM MWF values, although as expected, WM had significantly greater MWF values compared to cortical GM (Figure 6-G).

**Figure 6.**
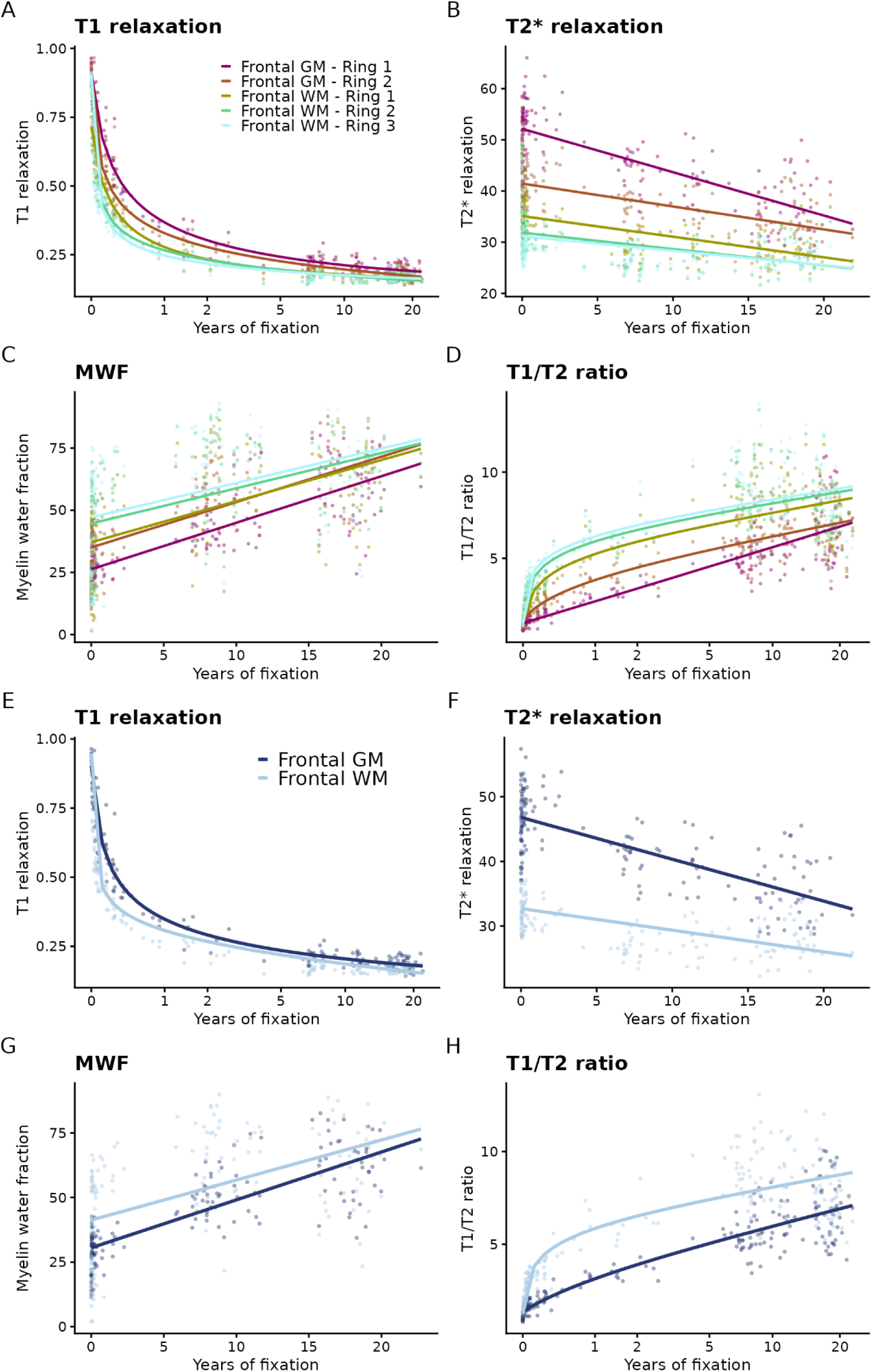
qMRI measurement changes associated with formaldehyde fixation at different tissue depths. A. T1 relaxation changes across rings. B. T2* relaxation changes across rings. C. MWF relaxation changes across rings. D. T1/T2 ratio changes across rings. E. T1 relaxation changes in cortical GM and WM. F. T2* relaxation changes in cortical GM and WM. G. MWF changes in cortical GM and WM.. H. T1/T2 ratio changes in cortical GM and WM. MWF: Myelin Water Fraction. GM: Cortical Gray Matter. WM: White Matter.

### Associations between qMRI measures, PMI, and pH

Linear regression models were used to examine the associations between PMI (Figure 7) and the qMRI measures, adjusting for the impact of formaldehyde fixation time based on the optimal model identified in the previous analyses. Compared to associations with formaldehyde fixation, PMI related changes were more limited. T1 relaxation times were not associated with PMI in cGM, WM, and most dGM and WMH regions. Only the hippocampus and frontal WMHs showed statistically significant increases with T1 relaxation time, after FDR correction. In contrast, T2* relaxation times increased with PMI across all cGM and WM regions, the hippocampus, thalamus, and frontal WMHs. Residualized T1/T2 ratio was only significantly associated with PMI in temporal WMHs. Finally, temporal and occipital cGM and hippocampal MWF significantly decreased with increase in PMI.

**Figure 7.**
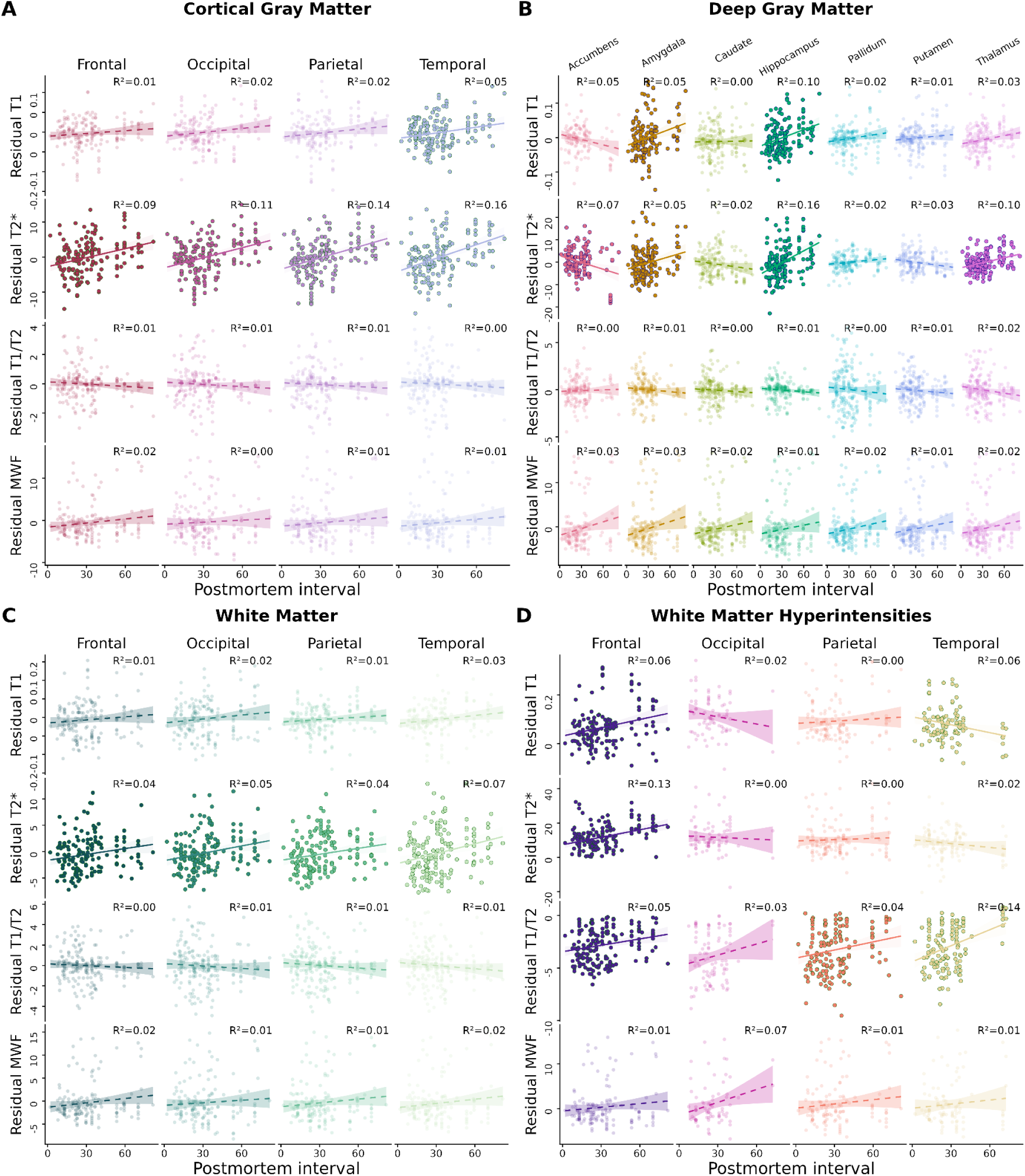
qMRI signal changes associated with postmortem interval (hours) accounting for the impact of formaldehyde fixation. A. Cortical gray matter. B. Deep gray matter. C. White matter. D. White matter hyperintensities.

Similar linear regression models were used to examine the associations between PMI and pH (Figure 8) and the qMRI measures, adjusting for the impact of formaldehyde fixation time based on the optimal model identified in the previous analyses. pH related changes were also limited, with residual T1 relaxation times showing a significant decline only in temporal cGM and residual T2* relaxation times showing a significant increase only in occipital cGM. Residual T1/T2 ratios significantly increased across all cGM regions, accumbens, and frontal, temporal, and parietal WMHs. Finally, MWF only showed significant associations with pH in occipital WMHs.

**Figure 8.**
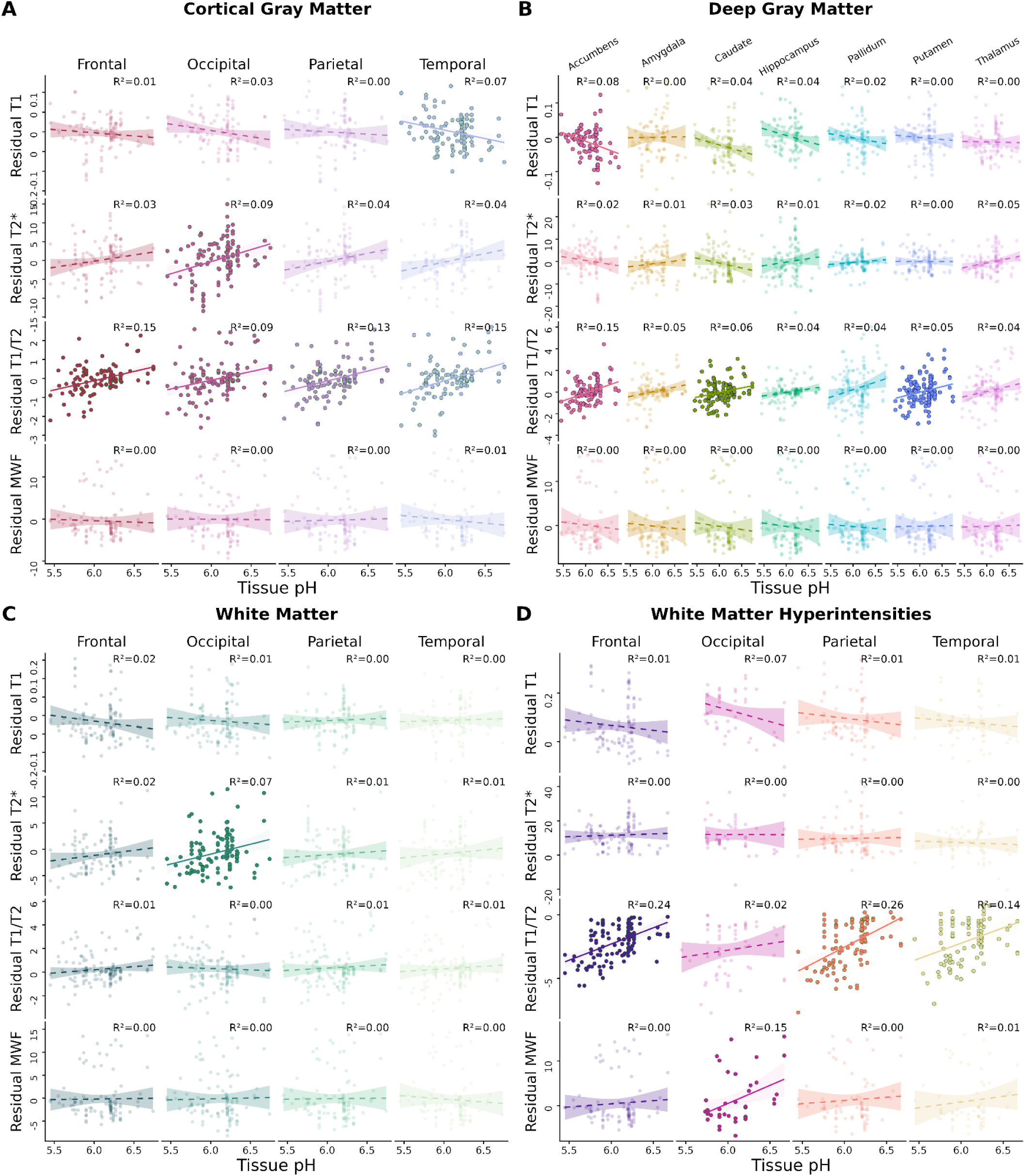
qMRI signal changes associated with pH accounting for the impact of formaldehyde fixation. A. Cortical gray matter. B. Deep gray matter. C. White matter. D. White matter hyperintensities.

## Discussion

The present study demonstrates and quantifies the effects of prolonged formalin fixation on various qMRI measurements (T1, T2, T2* relaxation times, and MWF) in a large and pathologically diverse sample of intact human brain specimens. Characterization of these effects is critical for studies that use postmortem MRI, whereby adequate adjustment for the impact of fixation is essential to derive qMRI based features that reflect biological tissue characteristics of interest such as presence of demyelination and neurodegenerative pathologies.

Formaldehyde fixation is a dynamic and nonlinear process (Raman et al., 2017). Formaldehyde forms covalent bonds between proteins in the tissue, maintaining the architecture of the cells. As the specimen is immersed in the fixative solution, formaldehyde diffuses throughout the tissue, reacting with the macromolecules to initiate protein cross-linking (Fox et al., 1985; Puchtler and Meloan, 1985). As such, fixation rate depends on the rate of exposure to the gradient of formalin, as well as other factors such as the PMI, pH, temperature, pressure, and uniformity of exposure, i.e. whether the tissue is immersed in a large container or exposure in certain areas are restricted (Thavarajah et al., 2012). Formaldehyde reaches the cGM first following immersion, resulting in more rapid formation of covalent cross-links and reduced initial degradation. As hemispheres are generally separated by a sagittal cut prior to immersion, formaldehyde can also reach dGM early in the fixation process, as the third ventricle is exposed. As fixation progresses, diffusion of the fixative may slow due to the presence of these newly formed bonds, which could explain the more gradual changes observed in GM compared to WM rings. Additionally, WM may begin to undergo degradation, such as delipidation, before formaldehyde penetrates the tissue. This could potentially account for the more rapid changes observed once formaldehyde eventually reaches the WM rings.

Both gray and white matter T1 and T2* relaxation times initially decreased with fixation. T1 values started at values similar to in vivo ranges (∼1 s) and rapidly declined reaching ∼0.5 s in the first few weeks to a year depending on the brain region, followed by stabilisation based on tissue type and proximity to the brain surface. T2 changes were slower and more subtle, and stabilisation was reached more gradually while preserving the initial regional contrasts for a long period of time. These results are consistent with prior literature (Birkl et al., 2018; Dawe et al., 2009; Pfefferbaum et al., 2004; Raman et al., 2017; Shatil et al., 2018; Tovi and Ericsson, 1992; Yong-Hing et al., 2005), and expand these findings past their relatively short assessment period (days to months). This is particularly important, as tissues stored at brain banks are commonly fixed for substantially longer windows, and quantifying the effect of fixation past a short few-month window will allow for use of these specimens in postmortem MRI studies, enabling direct comparison of qMRI and more traditional MRI signal with their gold standard cellular and pathological counterparts.

In contrast with T1 and T2* relaxation times, MWF and T1/T2 increased with prolonged fixation. Increase in MWF is associated with formaldehyde fixation in line with previous reports by Shatil and colleagues based on 2 postmortem specimens assessed from 0 to 43 days of fixation (Shatil et al., 2018), and expands these findings to a substantially longer postfixation period (i.e. 20 years). While the sensitivity of postmortem T1/T2 ratio to cortical demyelination has been previously studied (Zheng et al., 2022), to our knowledge the impact of formaldehyde fixation on T1/T2 ratio measures has not been previously examined. Our results show a rapid increase in T1/T2 ratio signals in the first few weeks (Figure 4) that stabilize but persist afterwards, and that these relationships differ across tissue types, with cGM regions showing a more linear pattern, while dGM and WM trajectories are more nonlinear and accelerated in the first few weeks of fixation.

Brain development follows an occipitofrontal (posterior-to-anterior) gradient of myelination (Barkovich et al., 1988; Boban et al., 2022; Yakovlev et al., 1967), resulting in substantial regional differences in myelin composition across brain regions. These differences are particularly pronounced in the frontal lobe, which is known to undergo myelin loss with aging (Boban et al., 2022). Previous work has suggested that formalin fixation may decompose the phospholipid structure of myelin, partially offsetting the overall reduction in relaxation times typically induced by fixation (Tovi and Ericsson, 1992). Similarly, differences in the T1 and T2* relaxation times obtained in the hippocampus and other regions may be consistent with the differential anterior-to-posterior changes in myelin content. Furthermore, prior studies by our team have examined the inversion of the GM-WM contrast in T1w images, reflecting the quality of the brain specimen fixation (Frigon et al., 2025, 2024). The GM-WM contrast inversion occurs first in the occipital lobe (Figure 1. Panel I), which experiences faster decreases in T1 relaxation times than other regions.

Additionally, we report interesting regional differences in fixation trajectories. T1 relaxation times decreased faster in certain deep GM regions compared to others. The hippocampus showed slower declines in T1 and T2* relaxation times, followed by the nucleus accumbens and the amygdala (Figures 2 and 3, Panel B). Compared to regions closer to the brain surface, dGM structures could be exposed to formaldehyde at lower rates when tissues are fixed by immersion as the fixative takes longer to penetrate in deep regions (McFadden et al., 2019). This may also be due to the presence of neuropathologies. The hippocampus is highly susceptible to hypoxic and/or ischemic injury, which has been linked to its sparser microvascular network (Johnson, 2023). The hippocampus is particularly susceptible to age-related neuropathologies including the accumulation of neurofibrillary tangles, amyloid plaques, TDP-43 proteinopathy or limbic predominant age-related TDP43 encephalopathy (LATE), as well as severe gliosis, neuronal atrophy or hippocampal sclerosis. The nucleus accumbens is closely connected to the hippocampus and affected by neuropathological lesions and the amygdala is a known epicenter for all these neuropathologies to co-occur in addition to Lewy-type α-sycucleinopathy and age-related tau astrogliopathy (ARTAG). In our cohort of specimens with neurodegenerative disorders, strong presence of neuropathological lesions might impact tissue integrity and penetration of formaldehyde in these regions. However, further investigations are necessary to fully understand the causes underlying the regional differences in qMRI metrics associated with formaldehyde fixation.

Due to its ability to generate robust segmentations from images with a range of intensity contrasts (Iglesias et al., 2023), SynthSeg was used to derive the initial tissue segmentation masks. This choice was particularly important for fresh to moderately fixed specimens (i.e. under 6 months), for which T1w tissue contrasts are more challenging to accurately delineate. Based on our experiments, SynthSeg was able to provide more robust tissue segmentations based on the T1w images compared to more recently published tools that have been developed for postmortem settings (Khandelwal et al., 2024). This is likely due to the substantial differences in fixation times and the fact that our T2w scans have a bright background, whereas the ex vivo tool was trained on Fomblin immersed specimens with a dark background.

Our study is not without limitations. Our sampling was more sparse for the 2.5-6 years fixation period due to restrictions in accepting donations during the COVID-19 pandemic. Due to acquisition time constraints, T2 relaxometry was performed based on the meGSE GRASE sequence. While this method has been well established and has previously been used for T2 quantification in postmortem settings (Piredda et al., 2021; Shatil et al., 2018), its relatively shorter TR introduces more T1-weighting than traditional T2-relaxometry methods. Furthermore, we have only examined the effect of 4% formaldehyde (i.e. 10% NBF) fixation as it is the most commonly used solution for long term preservation of postmortem human brain tissues. While this is the most common solution and concentration combination used in the brain banks, different concentrations of formaldehyde (e.g. 20% NBF) or differences in chemical composition of the fixative solutions can exert different effects on qMRI metrics or change the timescale of the observed effects (Frigon et al., 2025; Yong-Hing et al., 2005). Other parameters such as size of the brain, presence of pathologies, severity of atrophy, volume of fixative solution used can also impact the rates of fixation. As such, it is important to investigate the variability in qMRI metrics in large samples of specimens with different characteristics (e.g. age, postmortem interval, type and extent of neurodegenerative pathology). To our knowledge, our study is the first to systematically examine these effects in an unprecedented sample of aging brains with different neurodegenerative disorders.

Given that formaldehyde fixation also impacts the quality and staining intensity of histology and immunohistochemistry results (Frigon et al., 2026b; Ma et al., 2024; Pikkarainen et al., 2010; Webster et al., 2009), many studies opt for relatively short fixation periods (weeks to months) (McAleese et al., 2021, 2017, 2015; Zheng et al., 2022). Our results show that qMRI signals are particularly impacted during this period, and studies that aim to integrate qMRI measurements might benefit from choosing longer (e.g. 6 months to one year) and consistent fixation periods to maximize histology and immunohistochemistry results while avoiding significant fixation related variability in qMRI signals. Ex vivo MRI investigations of postmortem human brains with the goal of developing pathology specific MRI-based biomarkers have increasingly gained attention in recent years. Our findings provide valuable guidelines for future studies aiming to employ qMRI in postmortem settings. Based on these results, to minimize the effects of relaxation time variability caused by formaldehyde fixation, specimens can be reliably scanned past the 6-month fixation window whereby a linear adjustment would sufficiently account for the residual impact of formaldehyde fixation.

## Acknowledgements

We would like to acknowledge Dr. Gerald R. Moran (Head of Research, Innovation and Scientific Engagement, Siemens Healthcare Limited) who provided the GRASE sequence through a research application. We would also like to thank Danae Lussier-Dumouchel and Elena Drobotea for their help in MRI data acquisition. This project was supported by research funds from the Healthy Brains for Healthy Lives (HBHL), the Quebec BioImaging Network (QBIN), the Natural Sciences and Engineering Research Council of Canada (NSERC), the Canadian Institutes of Health Research (CIHR), Brain Canada, and ALS Canada-Brain Canada. Dadar also reports receiving research funding from the Fonds de recherche du Québec-Santé (FRQS, https://doi.org/10.69777/320107), CIHR, Alzheimer’s society research program (ASRP), and ALS Canada-Brain Canada. Zeighami reports funding from the FRQS (https://doi.org/10.69777/379799, and https://doi.org/10.69777/320107), CIHR, HBHL, NSERC, and ALS Canada-Brain Canada. The Douglas Brain Bank is supported by a platform support grant from Brain Canada. RM, WAG, and EMF are supported by scholarships and fellowships from the FRQS. CT is supported by a postdoctoral fellowship from the CIHR. ZA is supported by a doctoral NSERC Vanier scholarship.

